# Neuronal LRP4 directs the development, maturation, and cytoskeletal organization of peripheral synapses

**DOI:** 10.1101/2023.11.03.564481

**Authors:** Alison T. DePew, Joseph J. Bruckner, Kate M. O’Connor-Giles, Timothy J. Mosca

## Abstract

Synapse development requires multiple signaling pathways to accomplish the myriad of steps needed to ensure a successful connection. Transmembrane receptors on the cell surface are optimally positioned to facilitate communication between the synapse and the rest of the neuron and often function as synaptic organizers to synchronize downstream signaling events. One such organizer, the LDL receptor-related protein LRP4, is a cell surface receptor most well-studied postsynaptically at mammalian neuromuscular junctions. Recent work, however, has identified emerging roles for LRP4 as a presynaptic molecule, but how LRP4 acts as a presynaptic organizer, what roles LRP4 plays in organizing presynaptic biology, and the downstream mechanisms of LRP4 are not well understood. Here we show that LRP4 functions presynaptically at *Drosophila* neuromuscular synapses, acting in motor neurons to instruct multiple aspects of pre- and postsynaptic development. Loss of presynaptic LRP4 results in a range of developmental defects, impairing active zone organization, synapse growth, physiological function, microtubule organization, synaptic ultrastructure, and synapse maturation. We further demonstrate that LRP4 promotes most aspects of presynaptic development via a downstream SR-protein kinase, SRPK79D. SRPK79D overexpression suppresses synaptic defects associated with loss of *lrp4*. These data demonstrate a function for LRP4 as a peripheral synaptic organizer acting presynaptically, highlight a downstream mechanism conserved with its CNS function, and indicate previously unappreciated roles for LRP4 in cytoskeletal organization, synapse maturation, and active zone organization, underscoring its developmental importance.

## INTRODUCTION

The successful development of a synapse is an intricate process that requires coordinated steps mediated by diverse molecular players. Upon contact, pre- and postsynaptic partner neurons undergo steps of additional growth, transcriptional changes, structural and cytoskeletal remodeling, and recruitment of specialized synaptic proteins (Harris & Littleton, 2015). The complex developmental process at connections is crucial not only for the formation of the synapse but also its robust and reliable function. In the absence of normal synaptic development, neuronal communication is starkly impaired. Consequently, errors during synapse development can disrupt neuronal activity and underlie neurodevelopmental, neuropsychiatric, and even neurodegenerative disorders, including autism, epilepsy, and schizophrenia (Barnat et al., 2020; Bennett, 2011; Bonansco & Fuenzalida, 2016; Gilbert & Man, 2017; Grant, 2012; Melom & Littleton, 2011). Given the importance of the developmental precision required for synapse formation, there must be careful coordination across the synapse of multiple developmental steps – such coordination is often mediated by “synaptic organizer” proteins (Z. Li & Sheng, 2003; Scheiffele, 2003; Siddiqui & Craig, 2011). Understanding both the identity of, and the pathways by which, synaptic organizers coordinate aspects of synapse development is a critical step towards understanding both the development and dysregulation of the nervous system.

The low-density lipoprotein-related receptor LRP4 acts as a synaptic organizer at both invertebrate and vertebrate synapses (DePew & Mosca, 2021). As a cell surface receptor, LRP4 is optimally positioned to mediate signaling across the synapse during development and instruct downstream changes in the cell. In its most well-studied role in mammalian neuromuscular junction (NMJ) formation, LRP4 functions as a postsynaptic receptor for the synaptogenic ligand Agrin to initiate a cascade of postsynaptic events, beginning with phosphorylation of MuSK (Kim et al., 2008; Weatherbee et al., 2006; Zhang et al., 2008). MuSK phosphorylation leads to subsequent phosphorylation of Dok7 (Bergamin et al., 2010), followed by recruitment and clustering of postsynaptic proteins, including acetylcholine receptors (AchRs), Crk and Crk-L (Hallock et al., 2010), and activation of synapse-specific transcription (Burden et al., 2013). Prior to Agrin-binding, LRP4 instructs pre-patterning of AChRs during the initial steps of postsynaptic development, serving as a “stop” signal for the growth cone upon its arrival at the prospective synaptic region (Hata et al., 2018; Weatherbee et al., 2006; Yumoto et al., 2012). Consistent with such a role in shaping the projection, LRP4 also constrains MuSK-dependent synapse formation in zebrafish (Walker et al., 2021). Following synapse formation, LRP4 continues to function postsynaptically in NMJ maintenance, underlying its implication in disorders of the motor unit including amyotrophic lateral sclerosis (ALS) and myasthenia gravis (MG) (Barik et al., 2014; Pevzner et al., 2012; Rivner et al., 2017; Tzartos et al., 2014; Zhang et al., 2012). Further indicating its importance post-development, LRP4 also functions in the peripheral nervous system of zebrafish to promote regrowth of axons after injury (Gribble et al., 2018). In all, LRP4 functions in diverse, but primarily postsynaptic, roles to establish and maintain synaptic connections.

The potential presynaptic roles for LRP4 remain far less examined and understood. Phenotypic evidence following ablation of mouse LRP4 specifically in muscles demonstrates a role for muscle-derived LRP4 in presynaptic differentiation (Wu et al., 2012), which is suggested to occur via cleaved postsynaptic LRP4 acting on an unidentified presynaptic receptor (Yumoto et al., 2012). Further, the role of Agrin in either presynaptic function of LRP4 is not understood. As such, the identity of, and molecular mechanisms underlying, LRP4 function at peripheral presynapses both remain largely unknown. More is known about presynaptic LRP4 function in the central nervous system, however. In the *Drosophila* olfactory system, presynaptic LRP4 regulates active zone number and function in excitatory neurons via a downstream SR-protein kinase, SRPK79D. As *Drosophila* lack clear Agrin and MuSK homologues (Mosca et al., 2017), this suggests that fly LRP4 functions in the CNS independently from Agrin or MuSK. Despite this, how LRP4 in the fly acts at other stages of synaptic organization, development, and maturation, or how it may act transsynaptically are all unknown. A similar synaptogenic role for LRP4 exists in the mammalian central brain, where LRP4 regulates synapse number and dendrite morphology in an Agrin-dependent manner, though the contribution of MuSK in this pathway is unclear (Gomez et al., 2014; Handara et al., 2019; Karakatsani et al., 2017). Whether LRP4 function in the mammalian CNS is pre-or postsynaptic is not known, though some evidence suggests a postsynaptic etiology underlying these phenotypes (Tian et al., 2006). Taken together, while growing evidence implicates LRP4 as an important player in development of diverse synapses and through different pathways, including independently of Agrin and MuSK (DePew & Mosca, 2021), the precise mechanisms governing these roles remain unknown. Critical open questions remain regarding how LRP4 functions as a synaptic organizer, most notably what roles does presynaptic LRP4 play in influencing synaptic organization and what molecular mechanisms does LRP4 use independent from Agrin and MuSK to regulate synapse development?

The glutamatergic *Drosophila* larval neuromuscular junction (NMJ) is a powerful system to dissect the pre- and postsynaptic cellular mechanisms underlying synapse development (Figure 1A). Precise genetic tools and single-synapse imaging capability at the NMJ allow for careful investigation of synaptic development through cell-type specific analysis (Chou, Johnson, & Van Vactor, 2020; Collins & DiAntonio, 2007; Harris & Littleton, 2015; Keshishian et al., 2003). Moreover, there is a long history of translation of synaptic principles discovered at the *Drosophila* NMJ informing vertebrate work (Bellen et al., 2010, 2019; Jaiswal et al., 2012; Ma et al., 2022; Menon et al., 2013). Further, neurodevelopmental pathways of the fly peripheral nervous system have provided insight into neurodegenerative disorders due to the robust conservation between systems (Charng et al., 2014; Restrepo et al., 2022). NMJ development can be categorized by multiple overlapping stages, including synapse growth, cytoskeletal reorganization, active zone assembly, and maturation (Chou, Johnson, & Van Vactor, 2020). During synapse growth, addition of new synaptic boutons and branches occurs in an activity-dependent manner to accommodate the rapid growth of the larva (Ataman et al., 2008; Zito et al., 1999). As nascent boutons are added, microtubules are rearranged to accommodate new growth (Hummel et al., 2000; Roos et al., 2000). Development of the bouton further progresses through the addition of presynaptic release sites (Fouquet et al., 2009) precisely aligned across the synaptic cleft from postsynaptic receptor clusters (Schmid et al., 2006, 2008). Receptors are housed in a postsynaptic network of scaffolding and cytoskeletal proteins within the subsynaptic reticulum (SSR), a highly folded membranous structure surrounding each bouton (Guan et al., 1996; Pielage et al., 2006). As the synapse matures, additional receptors, scaffolding proteins, and cytoskeletal components are added to the postsynapse (Ataman et al., 2006; J. Li et al., 2007; Mathew et al., 2005; Mosca et al., 2012; Mosca & Schwarz, 2010; Owald et al., 2010; Packard et al., 2002, 2015; Restrepo et al., 2022; Speese et al., 2012). The precise pre- and postsynaptic roles of individual genes in each of these individual processes during development can be readily dissected at the NMJ, offering the opportunity to study how synaptic organizers like LRP4 influence diverse elements of development and their underlying cellular mechanisms. Overall, the *Drosophila* larval NMJ provides an excellent genetic model for investigating cellular mechanisms of development that can also be translated to the mammalian brain.

**Figure 1.**
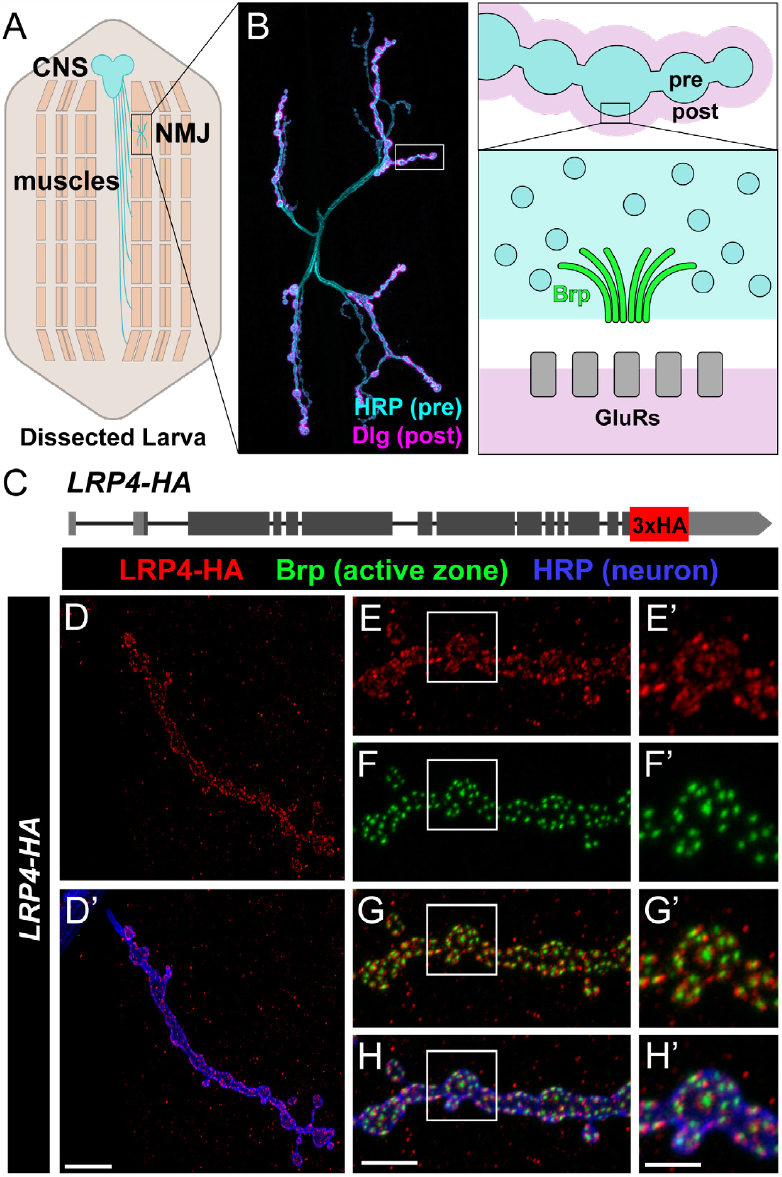
LRP localizes near active zones and is required for active zone organization. (**A**) Schematic of *Drosophila* third instar larva. Motor axons descend from the central nervous system (blue) to synapse with muscle fibers (tan). Muscles are organized in repeating segments along the length of the larva. (B) Representative confocal image (left) of a wild-type NMJ stained for HRP (cyan) to visualize neuronal membranes, and Dlg (magenta) to mark postsynapses. A diagram (right) depicting presynaptic boutons (blue) surrounded by the postsynaptic region (pink). Inset depicting the active zone (site of neurotransmitter release) and highlighting the active zone cytomatrix protein Bruchpilot (green). The active zone rests apposite a cluster of glutamate receptors (gray) located at the post-synaptic membrane. (**C**) Diagram of the *lrp4* genomic region indicating the locus of a 3xHA tag, inserted at the C-terminus. (**D**) Representative confocal image of an NMJ expressing endogenous LRP4-HA, stained with antibodies to HA (red) and HRP (blue). LRP4-HA expression is visible within the motor neuron. Scale bars = 10 μm. (**E-H**) Representative confocal image of an NMJ expressing endogenous LRP4-HA, stained with antibodies to HA (red), HRP (blue), and active zone marker Brp (green). Insets represent high magnification. LRP4-HA staining is visible in a punctate formation near active zones. Note the ring-like structure of HA staining surrounding Brp puncta. Scale bars = 5 μm, 2.5 μm (insets).

Despite roles in the CNS promoting axon targeting and olfactory function (Douthit et al., 2021; Mosca et al., 2017), how LRP4 regulates the multiple stages of synaptic development in the fly is not well understood. Critical open questions remain unanswered regarding LRP4 biology. Does LRP4 function presynaptically at peripheral synapses? What stages of synaptic development require LRP4? What cellular processes does LRP4 instruct during neurodevelopment? What are the downstream effectors of LRP4 in promoting synapse organization? To begin to answer these questions, we used the *Drosophila* NMJ as a model to study LRP4 function at the synapse. Here, we demonstrate that LRP4 is expressed presynaptically in motor neurons and localizes near active zones at the developing NMJ, presenting an opportunity to further investigate the cellular processes that presynaptic LRP4 regulates during development. Consistent with presynaptic expression, we find that LRP4 is required for nearly all aspects of synaptic development, including the organization of active zones, synaptic function, growth of the NMJ terminal, and organization of the microtubule cytoskeleton. In each case, LRP4 regulates synapse development by functioning cell-autonomously in presynaptic motor neurons. We also find that LRP4 is required for postsynaptic maturation, as loss of *lrp4* impairs recruitment of postsynaptic scaffolding proteins and formation of the postsynaptic spectrin cytoskeleton. LRP4 functions presynaptically to mediate its effects on synaptic maturation, suggesting both cell-autonomous and cell non-autonomous roles for LRP4. Finally, we demonstrate that LRP4 acts genetically via the downstream SR-protein kinase SRPK79D specifically in motor neurons to regulate development. Our work highlights the importance of LRP4 as a synaptic organizer that directs multiple cellular processes during development, and reveals novel roles for LRP4 in regulating microtubule organization and synapse maturation. LRP4 function through the kinase SRPK79D also highlights an Agrin / MuSK-independent mode of action. In all, these findings suggest shared presynaptic LRP4-dependent mechanisms between invertebrate and vertebrate synapse formation.

## RESULTS

### LRP4 is expressed presynaptically near active zones at the NMJ

In *Drosophila, lrp4* transcript expression is detected in motor neurons (Li et al., 2022) but this neither confirms protein expression nor provides the resolution needed to determine if LRP4 functions presynaptically at the NMJ or postsynaptically to interneuron populations. Thus, we first sought to determine if LRP4 protein is expressed in motor neurons and identify where LRP4 specifically localizes. We first confirmed LRP4 expression at the presynaptic NMJ by driving expression of GFP using an LRP4-GAL4 driver (Mosca et al., 2017; Pfeiffer et al., 2008) and observed GFP in motor neurons (Supplemental Figure 1A-B). To determine where LRP4 protein localizes in motor neurons, we used CRISPR-Cas9 genome editing (Gratz et al., 2013, 2014) to generate LRP4-HA, a fly line in which endogenous LRP4 is tagged with a 3x-HA epitope at the C-terminus (Figure 1C), as C-terminal tagging does not interfere with LRP4 function (Mosca et al., 2017). LRP4-HA flies were viable and showed no overt phenotypes, further indicating the tag did not interfere with normal function. We observed LRP4-HA staining in motor neurons (Figure 1D), supporting our GFP expression data. We also observed punctate LRP4-HA staining at NMJs within presynaptic boutons (Figure 1E-H). Specifically, co-staining with the active zone marker Bruchpilot (Brp) showed LRP4-HA localization near active zones, often organized in a punctate ring surrounding Brp puncta (Figure 1E-H). This indicates that LRP4 is localizes to presynaptic terminals, surrounding active zones. To further and independently confirm our findings, we also expressed a tagged *lrp4* transgene, UAS-LRP4-HA, in neurons via UAS/GAL4, and observed similar HA staining within boutons (Supplemental Figure 1C-D) and synaptic localization. Taken together, these data demonstrate that LRP4 protein is expressed in motor neurons and localizes presynaptically at the NMJ, suggesting that it may serve developmental and functional roles at or near the active zone.

### Perturbation of *lrp4* affects active zone organization and function

Active zones are comprised of a host of proteins surrounding a central cluster of calcium channels, allowing Ca^2+^-dependent and synchronous vesicle fusion and subsequent neurotransmitter release (Van Vactor & Sigrist, 2017). As we observe LRP4 presynaptically near active zones, we reasoned that LRP4 may be involved in active zone organization. To test our hypothesis, we disrupted LRP4 function using a previously generated null mutant (*lrp4*^*dalek*^), which lacks the entire coding region of *lrp4* (Mosca et al., 2017). At the NMJ, the active zone scaffolding protein Bruchpilot (the ortholog of vertebrate ELKS/CAST) regulates neurotransmission and active zone structure (Wagh et al., 2006) and is closely apposed to postsynaptic glutamate receptor tetramers containing the obligate GluRIIC / DGluRIII subunit (Collins & DiAntonio, 2007; Marrus et al., 2004) (Figure 1B). To first determine if LRP4 influences active zone and glutamate receptor cluster number, we stained for Bruchpilot and GluRIIC and quantified pre- and postsynaptic puncta (Figure 2A-B). In the absence of *lrp4*, we observed neither differences in the density of Brp (Figure 2C) or GluRIIC (Figure 2D) puncta nor in the ratio of Brp puncta to GluRIIC puncta (2E). We did, however, observe a significant increase in the number of unapposed active zones and postsynaptic receptor clusters per NMJ, defined as a Brp punctum lacking an apposite GluRIIC punctum or vice versa (Figure 2F). Unapposed GluRIIC puncta comprised 80% of apposition errors in the *lrp4* mutant (Supplemental Figure 2A-B). This increase in unapposed puncta suggests that, though *lrp4* does not affect active zone density, it is required for normal active zone apposition.

**Figure 2.**
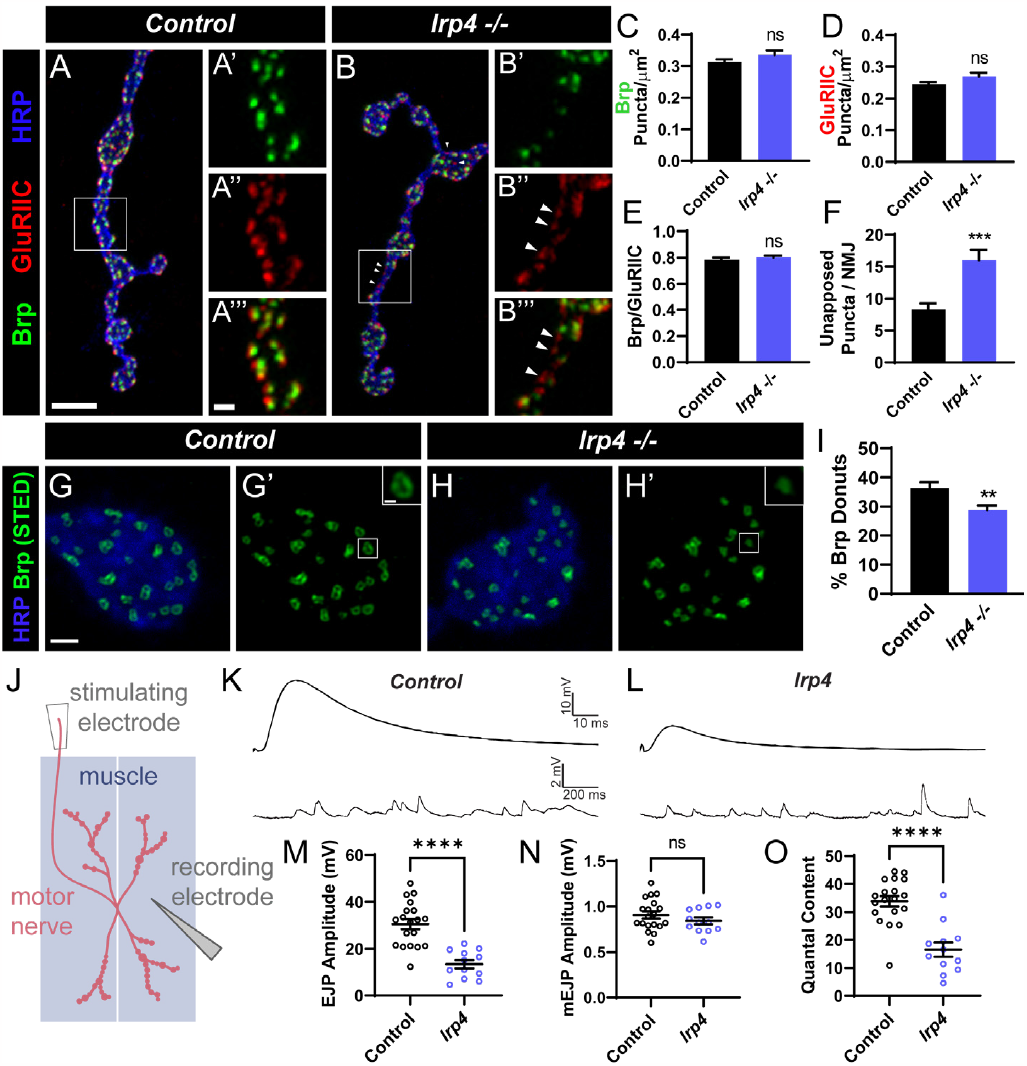
Loss of *lrp4* leads to defects in active zone apposition and function. (**A-B**) Representative confocal images of control (**A**) and *lrp4* mutant (**B**) NMJs stained with antibodies to Brp (green), GluRIIC (red), and HRP (blue). Arrowheads in insets indicate unapposed puncta. Scale bars = 5 μm, 2 μm (insets). (**C**) Quantification of Brp puncta density. (**D**) Quantification of GluRIIC density. (**E**) Quantification of the ratio of Brp to GluRIIC puncta. (**F**) Quantification of the number of unapposed puncta per NMJ. Unapposed puncta increase significantly following loss of *lrp4*. (**G-H**) Representative images of control (**G**) and *lrp4* mutant (**H**) NMJs stained with antibodies to HRP (blue) and Brp (green) visualized with STED microscopy. Insets show an example of a donut-shaped puncta. Scale bars = 1 μm, 200 nm (inset). (**I**) Quantification of percent of donut-shaped Brp puncta (with a center hole). The percent of donut-shaped puncta decreases following loss of *lrp4*. (**J**) Diagram of experimental setup for electrophysiological analyses. An electrode records from muscle fiber, either in the presence (to record EJPs) or absence (to record mEJPs) of external stimulation of the motor axon. (**K-L**) Representative EJP and mEJP traces recorded from wild-type (**H**) and *lrp4Del / y* (**I**) muscle. (**M**) Quantification of EJP amplitude. Loss of lrp4 results in an over 50% decrease in EJP amplitude. (**N**) Quantification of mEJP amplitude. mEJP amplitude is unchanged in *lrp4* mutants. (**O**) Quantification of quantal content. Quantal content is significantly decreased following loss of *lrp4*. For all experiments, data are shown as mean ± SEM. p** < 0.01, p*** < 0.001, p**** < 0.0001, ns = not significant. Significance was determined using a two-tailed Student’s t-test. *n* (**C-F**) ≥ 12 NMJs, 6 larvae. *n* (**G-H**) ≥ 47 boutons, 8 larvae. *n* (**M-O**) ≥ 12 NMJs, 5 larvae.

Though changes to active zone apposition were not accompanied by changes to active zone density, we could not rule out the possibility that LRP4 influenced the organization of individual active zones. With confocal imaging, active zones appear as individual puncta (Wagh et al., 2006) but when imaged using super-resolution STED microscopy, Brp puncta appear as donut-like structures in Type Ib boutons when oriented planar to the imaging axis (Fouquet et al., 2009; Jetti et al., 2023; Kittel et al., 2006). Defects in synaptic organization are often accompanied by changes to this donut structure, resulting in deformed active zones (Barber et al., 2018; Bruckner et al., 2017; Jetti et al., 2023; Liu et al., 2011). Using STED microscopy, we observed multiple Bruchpilot-positive active zone puncta with clear donut-shaped morphology in wild-type larvae (Figure 2G). We also observe a proportion of puncta in wild-type boutons that appear amorphous and lack a center hole; these likely represent donut-shaped puncta observed laterally (Figure 2G). In *lrp4* mutants, we observed significantly fewer puncta in Type Ib boutons that could be resolved as a donut and a concomitant increase in the proportion of amorphous active zone puncta (Figure 2H, I). Taken together, our data suggests that LRP4 is required both for normal apposition and organization of individual active zones.

To determine if the active zone defects we observed corresponded to functional deficits, we measured neurotransmission in NMJs lacking *lrp4*. We recorded both spontaneous and evoked potentials from muscles of wild-type and *lrp4* mutant larvae (Figure 2J-L). Loss of *lrp4* caused a 56% decrease in the amplitude of excitatory junctional potentials (EJPs), indicating impaired neurotransmission (Figure 2K-L, M). Interestingly, loss of *lrp4* did not affect the amplitude of spontaneous miniature EJP (mEJP) events (Figure 2K-L, N), suggesting that the defect observed could not solely be ascribed to a postsynaptic etiology. We calculated quantal content and determined that neurotransmitter release is significantly reduced in the absence of *lrp4* (Figure 2O). In all, our data indicates that LRP4 promotes the function, organization, and apposition of individual NMJ active zones, suggesting that LRP4 is essential for normal synaptic development.

### Neuronal LRP4 is critical for NMJ growth and microtubule organization

We next sought to determine if loss of *lrp4* affects NMJ synapse growth beyond its influence on active zone organization and function. We assessed overall NMJ growth by staining with antibodies to HRP to mark neuronal membranes (Figure 3A-B) and observed a 35% decrease in the number of synaptic boutons following loss of *lrp4* compared to wild-type (Figure 3I). To determine if this decrease in bouton number results from loss of *lrp4* in motor neurons, we performed tissue-specific rescue experiments in the *lrp4* mutant background and found that LRP4 expressed in motor neurons (Figure 3C), but not in muscles, is sufficient to rescue the observed bouton number phenotype (Figure 3I). Further consistent with LRP4 acting presynaptically, *lrp4* RNAi knockdown in motor neurons (Figure 3D), but not in muscle, recapitulates the reduction in bouton number (Figure 3I). Interestingly, overexpression of LRP4 in neurons increased bouton number (Supplemental Figure 3A-C), suggesting that LRP4 may act instructively in the motor neuron to control neuronal arborization and synapse formation. These data indicate that presynaptic LRP4 is required beyond active zones for normal NMJ growth.

**Figure 3.**
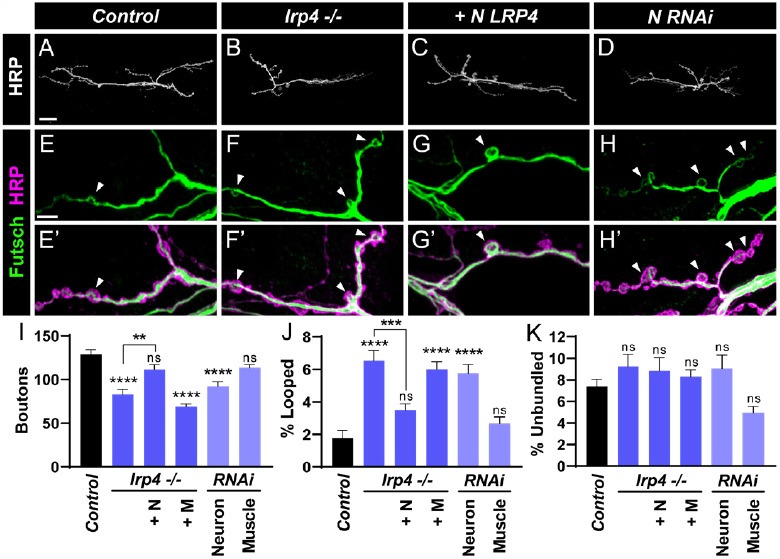
Motor neuron LRP4 is required for synapse growth and microtubule organization. (**A-D**) Representative confocal images of NMJs from control (**A**), *lrp4* mutant (**B**), *lrp4* mutant expressing LRP4 in motor neurons (**C**), and following knockdown of *lrp4* in motor neurons (**D**) stained with antibodies to HRP. Scale bars = 25 μm. (**E-H**) Representative confocal images of NMJs from control (**E**), *lrp4* mutant (**F**), *lrp4* mutant expressing LRP4 in motor neurons (**G**), and following knockdown of *lrp4* in motor neurons (**H**) stained with antibodies to Futsch (green) and HRP (magenta). Arrowheads indicate Futsch loops. Scale bars = 5 μm. (**I**) Quantification of bouton number, from experiments in (**A-D**); +N = expression in motor neurons, +M = expression in muscle. Loss of *lrp4* in motor neurons results in a significant decrease in bouton number. (**J**) Quantification of the percentage of boutons containing looped Futsch, from experiments in (**E-H**); +N = expression in motor neurons, +M = expression in muscle. Loss of *lrp4* in motor neurons significantly increased the percentage of boutons containing Futsch loops. (**K**) Quantification of the percentage of boutons containing unbundled Futsch; +N = expression in motor neurons, +M = expression in muscle. Loss of *lrp4* did not affect unbundled Futsch. For all experiments, data are shown as mean ± SEM. *p*\*\* < 0.01, *p*\*\*\* < 0.001, *p*\*\*\*\* < 0.0001, *ns* = not significant. Significance was calculated using one-way ANOVA, followed by Tukey’s test for multiple comparisons. n ≥ 13 NMJs, 8 larvae.

At the Drosophila NMJ, defects in synaptic growth are often associated with perturbations in microtubule cytoskeleton dynamics (Menon et al., 2013; Pennetta et al., 2002; Roos et al., 2000; Ruiz-Canada et al., 2004). The microtubule-associated protein Futsch / MAP1B colocalizes with microtubules specifically within motor neurons and is necessary for synaptic growth and stabilization of active zones (Hummel et al., 2000; Lepicard et al., 2014; Roos et al., 2000). As a potential mechanism underlying the disrupted growth in *lrp4* mutants, we further examined cytoskeletal organization using Futsch staining to visualize microtubule morphology (Figure 3E-H). In wild-type larvae, Futsch staining reveals branches of microtubules throughout the terminal, with looped microtubule or unbundled structures occasionally present within boutons. Loops are thought to indicate microtubule stability (Nechipurenko & Broihier, 2012; Roos et al., 2000; Shi et al., 2019), which may correlate with restricted growth (Chou, Johnson, Long, et al., 2020). Conversely, unbundled, diffuse Futsch is typically associated with disorganized microtubules (Roos et al., 2000). *lrp4* mutants show more than a three-fold increase in the percentage of boutons containing looped Futsch (Figure 3F, J), even despite a 35% reduction in the total number of boutons (Figure 3I). This increase in looped Futsch is suppressed by expression of LRP4 in motor neurons, but not in muscles (Figure 3G, J). Conversely, knockdown of *lrp4* in motor neurons increases the percentage of loops observed (Figure 3H), while knockdown of *lrp4* in muscle has no effect (Figure 3J). We also quantified the percentage of boutons containing unbundled Futsch and observed no significant differences across all examined genotypes (Figure 3K). The observed changes in Futsch indicate that presynaptic LRP4 regulates microtubule organization, and results in abnormally stabilized microtubules (but without affecting the unbundled population), which may contribute to the growth defects associated with *lrp4* loss.

### LRP4 in motor neurons is required for synapse maturation

Following the initial stages of growth, bouton addition, and active zone assembly, the synapse undergoes further steps to mature the nascent connection into a reliable synapse. During this process of synapse maturation, boutons recruit additional postsynaptic components, including glutamate receptors (Schmid et al., 2008), scaffolding proteins (Ataman et al., 2006; Packard et al., 2002), and cytoskeletal elements (Restrepo et al., 2022) to ensure the lasting strength of the synapse. Impaired synapse maturation can be observed as an increase in the number of immature boutons -termed “ghost boutons” -which are marked by presynaptic membrane but lack apposite postsynaptic markers like Dlg (Ataman et al., 2006; Piccioli & Littleton, 2014) and as a general reduction of the spectrin shell surrounding the postsynapse (Mosca & Schwarz, 2010; Restrepo et al., 2022).

We first examined if LRP4 promotes synaptic maturation by examining pre- and postsynaptic markers and quantifying the number of HRP-positive ghost boutons lacking the postsynaptic marker Dlg at *lrp4* mutant NMJs (Figure 4A-B). Compared to controls, loss of *lrp4* increased ghost boutons four-fold (Figure 4A-B, I). The defect in maturation can be rescued through expression of LRP4 in the motor neuron (Figure 4C), but not in muscle (Figure 4I), suggesting that presynaptic LRP4 is necessary for synaptic maturation and postsynaptic protein recruitment. Consistent with a presynaptic role for LRP4, presynaptic RNAi knockdown of lrp4 in motor neurons (Figure 4D), but not in postsynaptic muscles (Figure 4I), results in a significant increase in ghost boutons. We next examined postsynaptic β-spectrin staining surrounding boutons (Figure 4E-F). Loss of *lrp4* results in a 50% decrease in spectrin fluorescence intensity compared to controls (Figure 4E-F, J). This decrease can be rescued in the *lrp4* mutant background by expression of LRP4 in motor neurons (Figure 4G), but not in muscle (Figure 4J). Knockdown of *lrp4* in motor neurons (Figure 4D), but again not in muscles (Figure 4J), recapitulates the phenotype observed in *lrp4* null mutants. These data indicate that presynaptic LRP4 functions to ensure normal postsynaptic maturation. Further, we observe a requirement for LRP4 in cytoskeletal organization, here through regulation of the postsynaptic spectrin cytoskeleton, suggesting that presynaptic LRP4 may be broadly required for cytoskeletal organization on both sides of the synapse.

**Figure 4.**
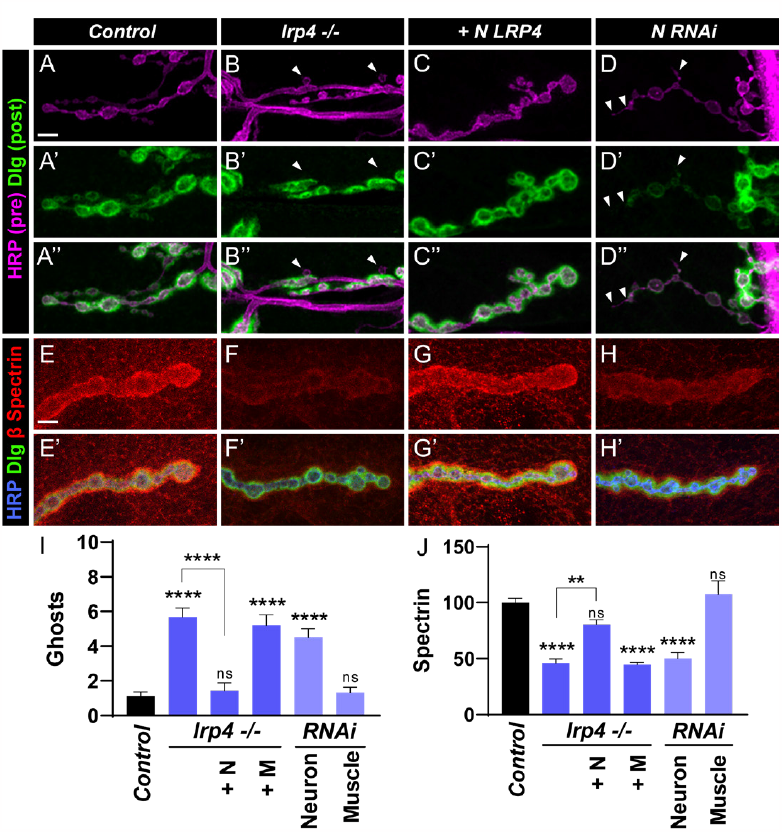
Motor neuron LRP4 is required for synapse maturation. (**A-D**) Representative confocal images of NMJs from control (**A**), *lrp4* mutant (**B**), *lrp4* mutant expressing LRP4 in motor neurons (**C**), and following knockdown of *lrp4* in motor neurons (**D**) stained with antibodies to Dlg (green) and HRP (magenta). Arrowheads indicate ghost boutons which lack Dlg staining. Scale bars = 5 μm. (**E-H**) Representative confocal images of NMJs from control (**E**), *lrp4* mutant (**F**), *lrp4* mutant expressing LRP4 in motor neurons (**G**), and following knockdown of *lrp4* in motor neurons (**H**) stained with antibodies to β-Spectrin (red), Dlg (green), and HRP (blue). Scale bars = 5 μm. (**I**) Quantification of ghost boutons, +N = expression in motor neurons, +M = expression in muscle. Loss of motor neuron LRP4 significantly increases the number of ghost boutons. (**J**) Quantification of spectrin fluorescence intensity levels (A.U.), +N = expression in motor neurons, +M = expression in muscle. Loss of LRP4 in motor neurons results in a significant decrease in spectrin fluorescence intensity levels at the NMJ. For all experiments, data are shown as mean ± SEM. *p*\*\* < 0.01, *p*\*\*\*\* < 0.0001, ns = not significant. Significance was calculated using one-way ANOVA,followed by Tukey’s test for multiple comparisons. n ≥ 14 NMJs, 7 larvae.

### Loss of *lrp4* perturbs synaptic ultrastructure

At the ultrastructural level, synaptic boutons appear as discrete structures that contain synaptic vesicles clustered around presynaptic active zones called T-bars (Feeney et al., 1998; Jia et al., 1993), and are surrounded by a membranous, folded structure, termed the subsynaptic reticulum (SSR). The SSR comprises the postsynaptic membrane, and contains neurotransmitter receptors, scaffolding proteins, and postsynaptic signaling machinery (Gan & Zhang, 2018; Guan et al., 1996; Lahey et al., 1994; Prokop, 2006) (Figure 5A). We sought to observe these structures at higher resolution, as structural defects in active zones and the SSR that are evident via electron microscopy (EM) are often not observed at the light level. We examined *lrp4* mutants using EM ultrastructural analysis to to better examine the effects of loss of function on synaptic organization. Compared to wild-type, we find that synaptic boutons in *lrp4* mutants have a 26% reduction in SSR area (Figure 5F) and a 19% reduction in SSR width (Figure 5G). The remaining SSR present in *lrp4* mutants has significantly reduced membrane infolding (Figure 5H); reduced SSR complexity is consistent with previous maturation mutants (Mosca et al., 2012; Mosca & Schwarz, 2010). Moreover, SSR complexity defects likely correspond to the decrease in spectrin levels we observe at the light microscopy level (Figure 4), as spectrin coincides with the SSR and is required for its formation (Pielage et al., 2006). We also observed regions of discontinuous or disorganized SSR in *lrp4* mutants that were never observed in wild-type boutons (Figure 5A-B). Taken together, these defects in the SSR suggest that LRP4 is required for the organization and normal biogenesis of postsynaptic membranes.

**Figure 5.**
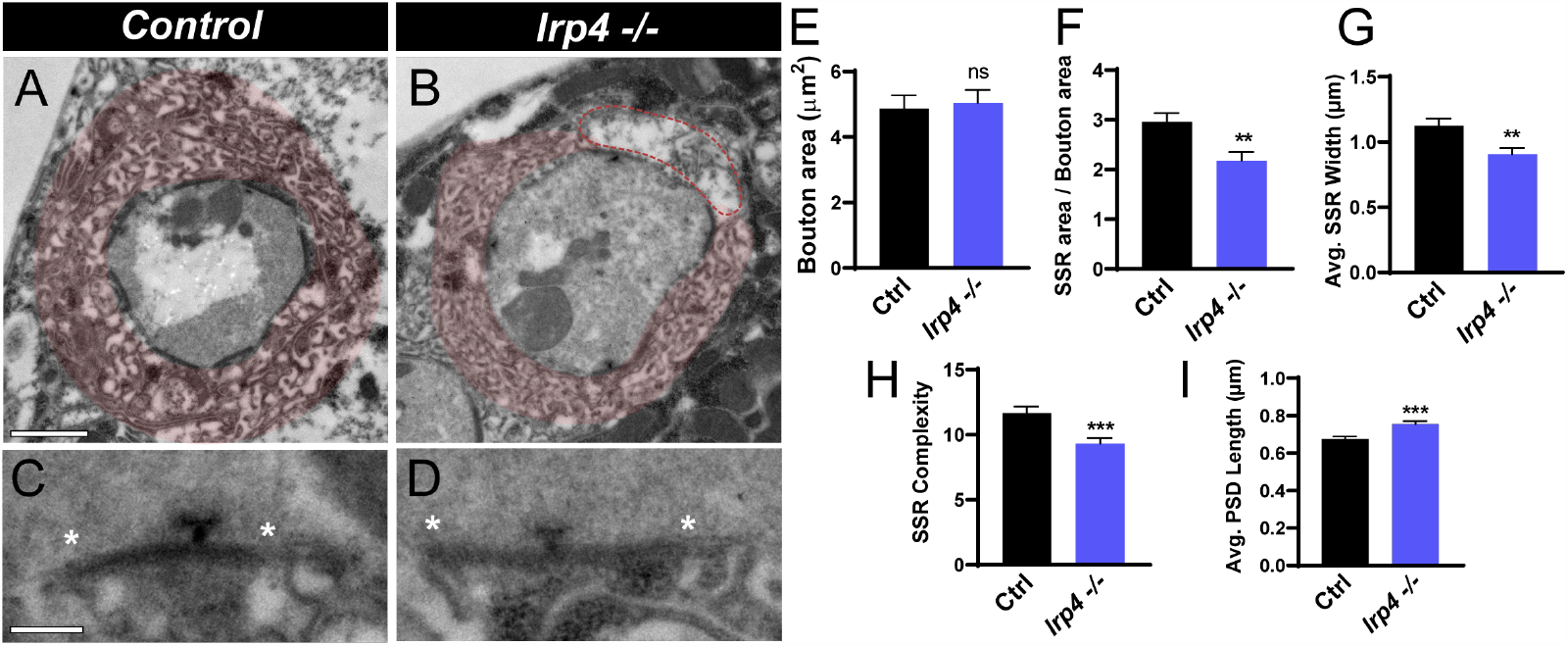
Loss of lrp4 leads to ultrastructural defects in membrane complexity. (**A-B**) Representative electron micrographs of boutons from control (**A**) and *lrp4* mutant (**B**) larvae, with SSR false-colored in red. Dotted lines in (**B**) indicate a region of disorganized SSR. Scale bars = 1 μm. (**C-D**) Representative electron micrographs of PSDs from control (**A**) and *lrp4* mutant (**B**) larvae. Asterisks denote the boundaries of PSDs. Scale bars = 200 nm. (**E**) Quantification of bouton area. Bouton area is unchanged in *lrp4* mutants. (**F**) Quantification of SSR area normalized to bouton area. SSR area is significantly decreased in *lrp4* mutants. (**G**) Quantification of average SSR width. SSR width is significantly decreased in *lrp4* mutants. (**H**) Quantification of average membrane crossings, a measure of SSR complexity. Loss of *lrp4* results in a significant decrease in membrane complexity. (**I**) Quantification of average PSD length. Average PSD length is decreased following loss of *lrp4*. For all experiments, data are shown as mean ± SEM. *p*\*\* < 0.01, *p*\*\*\* < 0.001, *ns* = not significant. Significance was determined using a two-tailed Student’s t-test. n ≥ 44 boutons, 3 larvae.

At the level of individual release sites, *lrp4* mutants displayed additional synaptic defects. Beyond synaptic maturation, the postsynaptic spectrin cytoskeleton is required for proper size of postsynaptic densities at the NMJ -loss of spectrin results in longer PSDs (Pielage et al., 2006). We quantified presynaptic parameters in our EM dataset and found that, though there were no changes evident in bouton perimeter or T-bar number / length (Supplemental Figure 4A-C), there was a significant increase in PSD length in *lrp4* mutants compared to wild-type (Figure 5I). Taken together with our SSR measurements, the data demonstrate an important role for LRP4 both in the biogenesis of synaptic membranes of the SSR and in development of the postsynaptic density, consistent with defects in both active zone organization and synaptic maturation.

### The SR protein kinase SRPK79D functions in the same genetic pathway as LRP4

Our data highlight a role for presynaptic LRP4 in multiple aspects of synaptic organization including bouton growth, microtubule organization, active zone apposition and structure, as well as synaptic maturation beyond previous understanding. We next sought to ascertain the downstream mechanism by which presynaptic LRP4 functions in promoting synaptic organization and to identify the specific downstream molecular effector required for LRP4 function. In mammalian systems, LRP4 often functions upstream of the kinase MuSK (Kim et al., 2008; Zhang et al., 2008) but as *Drosophila* lack a MuSK homologue, this kinase could not be the downstream effector for fly LRP4. Previous work in the *Drosophila* CNS implicated a different kinase in functioning downstream of LRP4 to regulate active zone number and olfactory function – the SR-protein kinase SRPK79D (Mosca et al., 2017). SR-family kinases were originally identified for their roles in mRNA splicing but were more recently found to function throughout the cell (Giannakouros et al., 2011), including in the nervous system (Arancibia et al., 2019; Bustos et al., 2020; Chan & Ye, 2013). At the *Drosophila* NMJ, SRPK79D localizes presynaptically (Supplemental Figure 5A) at active zones (Johnson et al., 2009), and regulates active zone assembly via Brp phosphorylation (Driller, Lutzkendorf, et al., 2019; Johnson et al., 2009; Nieratschker et al., 2009) but how SRPK79D is regulated at NMJ synapses and what molecular players function upstream of SRPK79D to influence synaptic organization, however, all remain unknown.

We hypothesized that SRPK79D may function downstream of LRP4 to regulate some or all aspects of LRP4-dependent synapse development. To test this hypothesis, we first assessed whether perturbation of *srpk79D* affects synapse development and growth. If SRPK79D functions downstream of LRP4 or otherwise in the same genetic pathway, we would expect the loss of *srpk79D* to phenocopy the loss of *lrp4*. To disrupt SRPK79D function, we used a previously validated *srpk79D* mutant, *srpk79D*^*atc*^ (Johnson et al., 2009). Loss of *srpk79D* significantly decreased bouton number (Figure 6A-B, Q) by 24%, similar to *lrp4* mutants, indicating a role for SRPK79D in NMJ growth. Beyond the similarity in bouton phenotype, we also observed similar alterations in the microtubule cytoskeleton. *srpk79D* mutants displayed an over three-fold increase in the percent of boutons containing Futsch loops (Figure 6E-F, R), indicating a shared phenotype with *lrp4* null mutations and demonstrating a role for SRPK79D in the organization of the microtubule cytoskeleton. Perturbations of *srpk79D* also showed defects in active zone organization and in synaptic maturation, similar to *lrp4* mutants. Loss of *srpk79D* resulted in more instances of unapposed Brp or GluRIIC puncta at active zones (Supplemental Figure 6A-C), a four-fold increase in the number of ghost boutons (Figure 6I-J, S), and a concomitant decrease in postsynaptic spectrin fluorescence (Figure 6M-N, T). These combined data suggest that, via multiple metrics and for distinct aspects of development, *srpk79D* mutants phenocopy the *lrp4* mutants and may share involvement in similar developmental events.

**Figure 6.**
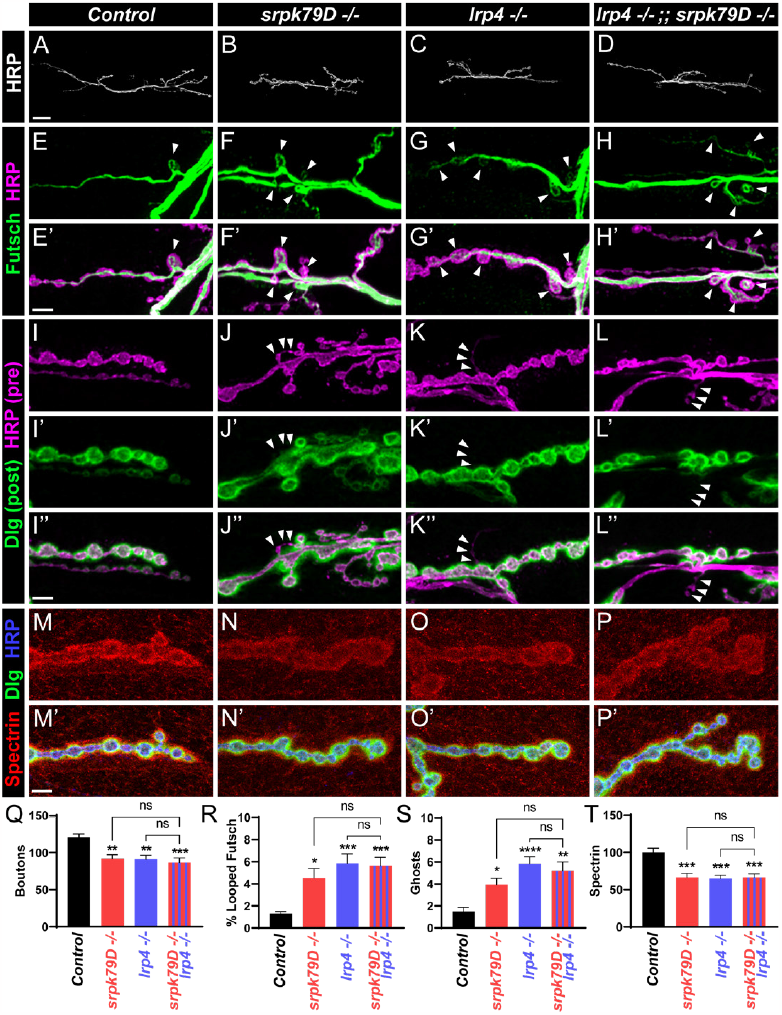
Loss of *srpk79D* phenocopies the loss of, and functions in the same genetic pathway as, *lrp4*. (**A-D**) Representative confocal images of NMJs from control (**A**), *srpk79D* mutant (**B**), *lrp4* mutant (**C**),and *lrp4, srpk79D* double mutant (**D**) larvae and stained with antibodies to HRP. Scale bars = 25 μm. (**E-H**) Representative confocal images of NMJs from control (**E**), *srpk79D* mutant (**F**), *lrp4* mutant (**G**), and *lrp4, srpk79D* double mutant (**H**) larvae and stained with antibodies to Futsch (green) and HRP (magenta). Arrowheads indicate Futsch loops. Scale bars = 5 μm. (**I-L**) Representative confocal images of NMJs from control (**I**), *srpk79D* mutant (**J**), *lrp4* mutant (**K**), and *lrp4, srpk79D* double mutant (**L**) larvae and stained with antibodies to Dlg (green) and HRP (magenta). Arrowheads indicate ghost boutons. Scale bars = 5 μm. (**M-P**) Representative confocal images of NMJs from control (**M**), *srpk79D* mutant (**N**), *lrp4* mutant (**O**), and *lrp4, srpk79D* double mutant (**P**) larvae and stained with antibodies to β-Spectrin (red), Dlg (green), and HRP (blue). Scale bars = 5 μm. (**Q**) Quantification of bouton number, from experiments in (**A-D**). (**R**) Quantification of percentage of boutons containing Futsch loops, from experiments in (**E-H**). (**S**) Quantification of ghost boutons per NMJ, from experiments in (**I-L**).(**T**) Quantification of spectrin fluorescence intensity levels (A.U.), from experiments in (**M-P**). In all cases, double mutation of *lrp4* and *srpk79D* does not result in phenotypes which differ significantly from mutation of either gene alone. For all experiments, data are shown as mean ± SEM. *p*\* < 0.05, *p*\*\* < 0.01, *p*\*\*\* < 0.001, *p*\*\*\*\* < 0.0001, ns = not significant. Significance was calculated using one-way ANOVA, followed by Tukey’s test for multiple comparisons. n ≥ 7 NMJs, 4 larvae.

The data next led us to determine if *lrp4* and *srpk79D* interact genetically and function together in the same pathway or in parallel pathways to instruct synapse development. We assessed a potential interaction using a double mutant approach; we reasoned that if LRP4 and SRPK79D function together, disruption of both would lead to similar phenotypes as mutation of either gene (i.e., would not enhance the phenotype). Conversely, if they functioned independently in parallel pathways, we would expect loss of both genes to enhance each other, resulting in a more severe phenotype. Importantly, we observed no significant difference in the double *lrp4*; *srpk79D* mutant compared to either of the single mutants in bouton number (Figure 6B-D, Q), microtubule organization (Figure 6F-H, R), ghost bouton number (Figure 6J-L, S), or spectrin fluorescence (Figure 6N-P, T). Our findings thus support a mechanism where LRP4 and SRPK79D likely function together in the same genetic pathway, and not in parallel pathways.

### SRPK79D functions downstream of LRP4 to instruct synapse growth, microtubule organization, and maturation

We finally sought to determine the epistatic relationship between *srpk79D* and *lrp4*; given the association of LRP4 with the synaptic membrane and the localization of SRPK79D at the synapse (Johnson et al., 2009), we reasoned that SRPK79D was most likely to function downstream of LRP4 (Mosca et al. 2017). We hypothesized that if SRPK79D functions downstream of LRP4, then overexpressing SRPK79D in the *lrp4* mutant background would be sufficient to suppress the phenotypes associated with *lrp4* loss. Moreover, given that our data indicates that LRP4 functions presynaptically, SRPK79D should also function in the motor neuron to regulate synaptic organization. To test this hypothesis, we expressed a venus-tagged SRPK79D in motor neurons of *lrp4* mutants and compared these to *lrp4* mutants similarly expressing a control GFP transgene in motor neurons (Figure 7). To quantify NMJ growth, we counted synaptic boutons (Figure 7A-C) and observed suppression of the decreased bouton number phenotype following overexpression of SRPK79D (Figure 7M). To assess microtubule organization, we assessed the percentage of boutons containing looped Futsch (Figure 7D-F), and again observed suppression of the *lrp4* mutant phenotype with overexpression of SRPK79D (Figure 7N). As metrics of synapse maturation, we assessed ghost boutons (Figure 7G-I, O) and spectrin fluorescence intensity (Figure 7J-L, P), and in both cases observed suppression of lrp4 mutant phenotypes following motor neuron overexpression of SRPK79D. To ensure that overexpression of SRPK79D alone does not cause any confounding phenotypes, we overexpressed SRPK79D in motor neurons in a wild-type background and observed no significant changes in multiple metrics of synaptic organization compared to control (Supplemental Figure 7). These data indicate that SRPK79D overexpression is sufficient to suppress *lrp4* mutant phenotypes in synapse growth, cytoskeletal organization, and synapse maturation. This suggests that SRPK79D functions downstream of presynaptic LRP4 in motor neurons to regulate diverse developmental processes and establishes a core neuronal signaling pathway that promotes pre- and postsynaptic NMJ development.

**Figure 7.**
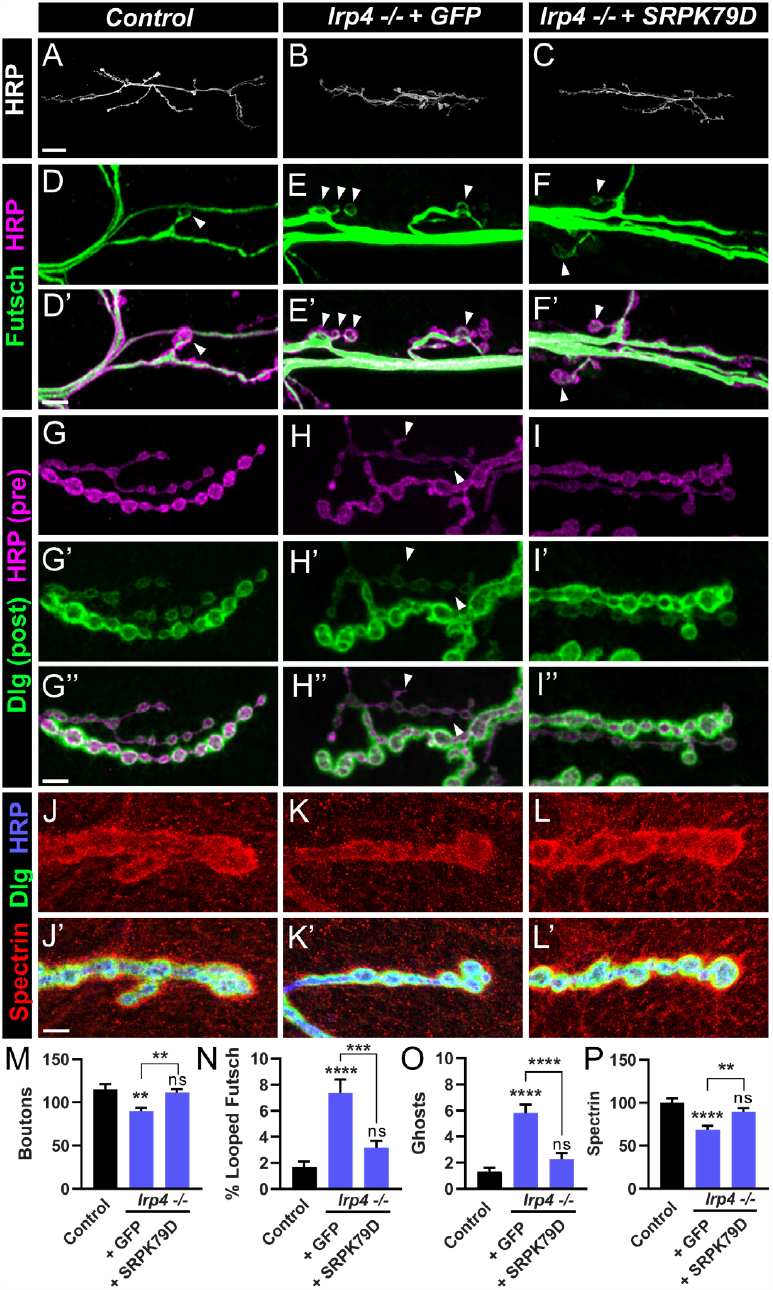
Overexpression of SRPK79D suppresses *lrp4* mutant phenotypes. (**A-C**) Representative confocal images from control (**A**), and *lrp4* mutant NMJs expressing GFP (**B**) or SRPK79D (**C**) in motor neurons, stained with antibodies to HRP. Scale bars = 25 μm. (**D-F**) Representative confocal images from control (**D**), and *lrp4* mutant NMJs expressing GFP (**E**) or SRPK79D (**F**) in motor neurons, stained with antibodies to Futsch (green) and HRP (magenta). Arrowheads indicate Futsch loops. Scale bars = 5 μm. (**G-I**) Representative confocal images from control (**G**), and *lrp4* mutant NMJs expressing GFP (**H**) or SRPK79D (**I**) in motor neurons, stained with antibodies to Dlg (green) and HRP (magenta). Arrowheads indicate ghost boutons. Scale bars = 5 μm. (**J-L**) Representative confocal images of control (**J**), and *lrp4* mutant NMJs expressing GFP (**K**) or SRPK79D (**L**) in motor neurons, stained with antibodies to β-Spectrin (red), Dlg (green), and HRP (blue). Scale bars = 5 μm. (**M**) Quantification of bouton number, from experiments in (**A-C**). (**N**) Quantification of the percentage of boutons containing Futsch loops, from experiments in (**D-F**). (**O**) Quantification of ghost boutons per NMJ, from experiments in (**G-I**). (**P**) Quantification of spectrin fluorescence intensity levels (A.U.), from experiments in (**J-L**). In all cases, overexpression of SRPK79D in the *lrp4* mutant background results in rescue of the phenotype. For all experiments, data are shown as mean ± SEM. *p*\*\* < 0.01, *p*\*\*\* < 0.001, *p*\*\*\*\* < 0.0001, *ns* = not significant. Significance was calculated using one-way ANOVA, followed by Tukey’s test for multiple comparisons. n ≥ 7 NMJs, 4 larvae.

## DISCUSSION

The coordination of intricate cellular processes during synapse development is critical to successfully form robust and lasting connections. Here we find that the cell surface receptor LRP4 acts presynaptically at peripheral synapses as a master organizer of multiple elements of synapse development, including growth, cytoskeletal structure, active zone organization, and synapse maturation. We also show that the SR-protein kinase SRPK79D functions downstream of LRP4 at the NMJ to promote these elements of synaptic development. These findings begin to answer critical open questions in LRP4-related and synaptic biology. Though postsynaptic functions of LRP4 are well documented (DePew and Mosca, 2021), how LRP4 may act presynaptically is notably less well understood. First, we highlight that LRP4 functions in presynaptic motor neurons at neuromuscular synapses which has previously been unclear and controversial. Second, we reveal that LRP4 influences a range of synaptic developmental processes leading to the development of a reliable connection. Third, we identify novel roles for LRP4 in regulating multiple aspects of the synaptic cytoskeleton and in synaptic maturation. Finally, we identify a shared downstream mechanism for LRP4 in the SRPK79D kinase that unites the roles of LRP4 as a master regulator of synaptic organization. These findings not only contribute to our understanding of mechanisms of synapse development in Drosophila but also inform our understanding of LRP4 function at other synapses. Given the mechanistic conservation between central and peripheral synapses in Drosophila (Mosca et al. 2017), our work provides unique insight on potential mechanisms in the mammalian CNS, where the precise developmental mechanisms LRP4 employs remain elusive (Gomez et al., 2014; Handara et al., 2019; Karakatsani et al., 2017).

### LRP4 as a presynaptic organizer of active zones

Pioneering work in the last 20 years identified a crucial role for LRP4 in mammalian NMJ synaptogenesis, stationed at the postsynaptic membrane and serving as the coreceptor for the synaptogenic ligand Agrin (Kim et al., 2008; Weatherbee et al., 2006; Zhang et al., 2008). There, muscle-derived LRP4 is essential to instruct various steps of postsynaptic development. LRP4 also plays an important role in presynaptic differentiation, as LRP4 is likely cleaved and serves as a retrograde signal to instruct motor neuron development (Yumoto et al., 2012). In this last case, the source of LRP4 is still postsynaptic but functions transsynaptically. Other presynaptic roles for LRP4 at the NMJ, however, have remained controversial. Some evidence suggests that LRP4 acts presynaptically in synaptic maintenance (Wu et al., 2012) but this is not well understood and the image of LRP4 has remained as a solely postsynaptic protein. Recent work in both vertebrate and invertebrate systems has begun to suggest a duality for LRP4, as both a presynaptic and a postsynaptic factor. In the central nervous system, evidence highlights a role for LRP4 in regulating synapse formation (Gomez et al., 2014; Handara et al., 2019; Karakatsani et al., 2017; Mosca et al., 2017). LRP4 acts presynaptically at *Drosophila* olfactory synapses to regulate active zones, though the mechanistic insight beyond active zones in other developmental processes is not yet understood and any potential postsynaptic role has not been identified (Mosca et al., 2017). At mammalian central synapses, LRP4 is enriched at pre- and postsynaptic membranes and its perturbation leads to both pre- and postsynaptic defects (Karakatsani et al., 2017) but the underlying mechanisms are also not understood. In the mammalian brain, moreover, where LRP4 functions to regulate synaptic biology is unknown, though some evidence suggests it may act postsynaptically (Tian et al. 2006). Using the fly NMJ as a model synapse, we sought to address potential roles of LRP4 with cell-type specific precision. We find that LRP4 is expressed presynaptically in motor neurons (Figure 1), presenting an opportunity to study its presynaptic roles during multiple aspects of development at a glutamatergic synapse. Acting presynaptically, we find that LRP4 controls active zone apposition and structure, synaptic function, microtubule organization, and bouton growth. Further, we observe that LRP4 localizes to the periactive zone, consistent with its localization at *Drosophila* central synapses colocalized with, or adjacent to, active zones (Mosca et al., 2017). Intriguingly, we find that presynaptic LRP4 influences aspects of postsynaptic maturation, highlighting not just autonomous roles for LRP4 in presynaptic biology but also a transsynaptic function for presynaptic LRP4 in regulating postsynaptic protein recruitment that promotes maturation. How these roles may be conserved in mammalian systems, especially in the CNS, where the precise function of LRP4 is not fully understood, remains a fascinating topic for future study and offers critical insight into the presynaptic functions of LRP4.

### LRP4 as a cytoskeletal regulator during growth and maturation

The importance of cytoskeletal dynamics in regulating neuronal, axonal, and synaptic biology cannot be understated (Goellner & Aberle, 2012; Lasser et al., 2018; Poulain & Sobel, 2010) but how cytoskeletal activity is integrated with, and responds to, synaptic signals from the cell surface is less well understood. Indeed, how LRP4 influences the morphological changes associated with growth or development in any system remains unknown. Our discovery of a role for presynaptic LRP4 in synaptic growth (as measured by bouton number) and subsequently, cytoskeletal organization, provides a unique opportunity to associate synaptic organizers at the membrane with the cytoskeleton. Microtubule dynamics greatly influence synaptic growth in multiple systems, and in Drosophila, perturbation of microtubule regulators can result in dramatic changes in bouton number (Chou, Johnson, Long, et al., 2020; Lepicard et al., 2014; Mosca et al., 2012; Roos et al., 2000; Ruiz-Canada et al., 2004; Shi et al., 2019). Consistent with this idea, we found, in the absence of presynaptic LRP4, an increase in looped microtubule structures, suggesting altered cytoskeletal stability. Improper microtubule stabilization can result in reduced plasticity of the cytoskeleton and may correlate with decreased growth (Chou, Johnson, & Van Vactor, 2020), as well as neuronal branching and synaptic morphology (Parato & Bartolini, 2021). We propose that presynaptic microtubules are improperly stabilized in the absence of LRP4, leading to less synaptic growth. As components of the microtubule cytoskeleton also contribute to the organization and stability of active zones (Koch et al., 2008; Lepicard et al., 2014), it remains a possibility that the defects in active zone organization we observe also result from the contribution of LRP4 to microtubule organization. Interestingly, previous work suggests that Futsch / MAP1B functions, in part, as an intermediate between active zones and microtubules, potentially contributing to active zone stability (Lepicard et al., 2014). As a result, we suggest that altered microtubule stability may signify the primary defect in lrp4 mutants from which alterations in bouton growth and active zone organization stem. As LRP4 localizes near active zones, it may thus serve as a nexus between the active zone machinery and Futsch, working to stabilize active zones and enable proper growth.

The regulation of cytoskeletal stability by LRP4 posits an intriguing hypothesis for mechanistic conservation at vertebrate synapses. Mammalian LRP4 in CNS neurons is essential for dendritic arborization and morphogenesis (Handara et al., 2019; Karakatsani et al., 2017). As in *Drosophila*, mammalian dendritic growth and branching (Poulain & Sobel, 2010), and synapse formation (Parato & Bartolini, 2021) rely on rearrangement of microtubules during development. This similarity raises the tantalizing possibility that LRP4 may also function in the mammalian CNS to instruct microtubule organization, thus contributing to synapse and neurite development. Given the importance of microtubule organization for diverse aspects of synapse function and the contribution of microtubule dynamics to neurodevelopmental disease (Lasser et al., 2018), this insight into a potentially conserved role for LRP4 in microtubule dynamics may inform our understanding of synapse development in both health and disease.

Cytoskeletal organization also plays an important role in synaptic maturation. Postsynaptic maturation involves the recruitment of postsynaptic components including the spectrin cytoskeleton to ensure that nascent boutons become functional, lasting connections (Ataman et al., 2006; J. Li et al., 2007; Mathew et al., 2005; Mosca et al., 2012; Mosca & Schwarz, 2010; Owald et al., 2010; Packard et al., 2002, 2015; Restrepo et al., 2022; Speese et al., 2012). Presynaptic LRP4 is essential for postsynaptic spectrin recruitment and SSR organization, indicating first, a transsynaptic role for LRP4 signaling and second, a deeper role in organizing the cytoskeleton. The picture that emerges from this work is that presynaptic LRP4 is essential to regulate the cytoskeletal elements that promote development at multiple stages. This markedly contributes to our growing understanding of synapse maturation mechanisms. In *Drosophila*, many pathways influence synapse maturation (Mosca et al., 2012; Owald et al., 2010; Packard et al., 2015; Sulkowski et al., 2016), including Wnt signaling (Ataman et al., 2006; Mosca & Schwarz, 2010; Restrepo et al., 2022). Binding of Wg, the *Drosophila* Wnt, to Frizzled receptors initiates a range of downstream signaling pathways, including formation of the postsynaptic density and SSR, and perturbation of this pathway results in an increase in ghost boutons and a decrease in spectrin levels (Ataman et al., 2006; Mosca & Schwarz, 2010; Restrepo et al., 2022). Further, Wnt signaling is involved in remodeling of the microtubule cytoskeleton (Gögel et al., 2006; Packard et al., 2002). Interestingly, another member of the LRP family, LRP5/6 (*Drosophila* Arrow), serves as the Frizzled coreceptor in Wnt signaling pathways at the *Drosophila* NMJ (Miech et al., 2008; Wehrli et al., 2000). These data suggest the possibility that LRP4, like LRP5/6, might also function to regulate Wnt signaling. While these pathways share similarities, there are notable differences which suggest LRP4 does not simply function to promote Wnt signaling. If this were the case, we would expect that loss of *lrp4* would phenocopy loss of *wg*, and decrease, rather than increase, Futsch loops (Packard et al., 2002). This does not rule out the possibility, however, that LRP4 may function with Wnt pathways, perhaps through a more complicated regulatory mechanism. It does suggest, however, that LRP4 acts via some (if not all) Wnt-independent pathways. Drawing mechanistic evidence from elsewhere in development, including in mammalian early forebrain development, LRP4 functions as an inhibitor of Wnt signaling pathways (Ahn et al., 2013, 2017; Choi et al., 2009; Geng et al., 2023). These data suggest that, at the *Drosophila* NMJ, LRP4 may be involved in the complex regulation of Wnt signaling, perhaps functioning as an inhibitor of Wnt pathways.

### SRPK79D functions with LRP4 to instruct development

SR-protein specific kinases (SRPKs) were initially discovered to phosphorylate SR-proteins and promote the subsequent nuclear import of mRNA splicing factors (Zhou & Fu, 2013). More recent work, however, has demonstrated numerous roles in the nervous system, including in development and disease (Bustos et al., 2020; Chan & Ye, 2013; Giannakouros et al., 2011; Hogg & Findlay, 2023). One *Drosophila* SRPK homologue, SRPK79D, has emerged as an important player in active zone assembly at the NMJ (Johnson et al., 2009; Nieratschker et al., 2009). In the absence of SRPK79D, active zone components aggregate prior to deposition at synaptic contacts, leading to reduced synaptic function, improper axonal localization of active zone material, and smaller release sites (Johnson et al., 2009; Nieratschker et al., 2009). To date, however, no clear upstream interactor has been identified for SRPK79D at the NMJ, leaving its regulatory mechanisms undiscovered. We found that not only do *srpk79D* mutants phenocopy *lrp4* mutants in multiple parameters of synaptic organization, they act in the same genetic pathway. Further, SRPK79D overexpression can rescue the phenotypes of *lrp4* mutants, suggesting that SRPK79D functions downstream of LRP4, revealing a regulatory mechanism for SRPK79D. We posit a model wherein LRP4 signals from the cell membrane through SRPK79D to influence development through regulation of the cytoskeleton. SRPK79D modulates assembly of active zones via phosphorylation of Bruchpilot, presenting a parsimonious route by which this pathway can influence active zone organization (Driller, Lutzkendorf, et al., 2019). Thus, our identification of LRP4 as the first upstream modulator of SRPK79D places it within a cascade where Bruchpilot functions downstream. An orthologous mammalian SR-protein kinase, SRPK2, also functions in assembling active zones, indicating conservation of this presynaptic developmental mechanism (Arancibia et al., 2019).

Beyond active zone assembly, we find that LRP4 and SRPK79D are required for pre- and postsynaptic organization of cytoskeletal components. How LRP4 and SRPK79D might influence cytoskeletal organization remains an open question. One possibility arises from recent work on mammalian SRPK2 demonstrating a role for the kinase in suppressing microtubule polymerization in neurons. SRPK2 exerts its function in microtubule polymerization through phosphorylation of the microtubule associated protein Tau (Hong et al., 2012). Like Futsch, Tau is involved in microtubule stabilization (Götz et al., 2006; G. Lee & Rook, 1992), raising the possibility that mammalian and *Drosophila* SRPKs may function in the nervous system through similar mechanisms to regulate the delicate balance of microtubule organization. In such a pathway, SRPK79D may influence the Futsch phosphorylation and thus regulate microtubule dynamics at the NMJ. That potential mechanism for SRPK79D has interesting parallels to that of another kinase, Shaggy (GSK3β), which influences microtubule stability and synapse growth by phosphorylating Futsch (Franco et al., 2004; Gögel et al., 2006). While loss of Shaggy increases microtubule loops, it also increases bouton number (Franco et al., 2004), in contrast to the disruption of *lrp4* / *srpk79D*. This suggests that multiple concurrent mechanisms regulate the complex balance of microtubule dynamics. It is possible that differences in temporal regulation, where increased stability earlier or later in development may influence growth differently. Effectors of microtubule stability may also exert their influence through different mechanisms, which consequently have differing effects on growth. Understanding the intersection of multiple mechanisms involving microtubule stability and phosphorylation targets will provide insight into the complex cytoskeletal dynamics that underly synapse development. Further, whether LRP4 and an SR-protein kinase like SRPK2 (Arancibia et al., 2019; Hong et al., 2012) may also function together in mammalian neurons to instruct cytoskeletal organization and influence neurite growth, active zone assembly, and maturation remains an intriguing possibility.

### Limitations of this study

Our analysis of LRP4 in active zone assembly demonstrates its diverse roles in synapse biology. The importance of LRP4 in assembling active zones is conserved across systems – in the *Drosophila* brain, LRP4 is required for proper synapse number (Mosca et al., 2017). Likewise, in mammals, loss of LRP4 at central synapses results in decreased synapse density (Handara et al., 2019; Karakatsani et al., 2017). Although we observed no changes in active zone density at the NMJ in *lrp4* mutants, the total number of active zones is likely reduced since fewer boutons are present, demonstrating a common role for LRP4 across systems and species in promoting synapse number. Further, while we observe a significant decrease in neurotransmitter release in *lrp4* mutants, the defects in active zone organization and apposition appear comparatively mild. How the organizational defects we observe in active zone structure, cytoskeletal organization, or synaptic apposition may account for this functional deficit remains unknown. One possibility is that LRP4 functions in another aspect of presynaptic development, aside from those elucidated here. Recent work in the mammalian central nervous system presents another interesting possibility. There, astrocytic LRP4 is implicated in the modulation of glutamate release (Sun et al., 2016). This remains an important future direction, though it is unlikely that glial LRP4 plays a significant role in the aspects of development outlined here, as we observe rescue of *lrp4* mutant phenotypes following expression of LRP4 in motor neurons, and *lrp4* RNAi in neurons recapitulates whole-animal mutants. However, a glial role for LRP4 in Drosophila, perhaps in modulating glutamate release in the CNS, remains a fascinating topic for future investigation.

**ACKNOWLEDGMENTS**

We thank Michael Aimino, Kristen Davis, Michael Parisi, and S. Zosimus for their comments on the manuscript. We deeply appreciate the gifts of reagents from Graeme Davis, Aaron DiAntonio, and Ron Dubreuil. We acknowledge the Bloomington Drosophila Stock Center (NIHP40OD018537) for providing stocks used in this study. Monoclonal antibodies used in this study were obtained from the Developmental Studies Hybridoma Bank, created by the National Institute of Child Health and Human Development (NICHD) of the NIH and maintained at the University of Iowa Department of Biology, Iowa City, IA. We also acknowledge the 2D & 3D Electron Microscopy Laboratories Shared Service Center at Thomas Jefferson University for EM support, and the Bioimaging Shared Resource of the Sidney Kimmel Cancer Center (NCI 5 P30 CA-56036), especially Jason Hill, for assistance and training in STED imaging and deconvolution.

## MATERIALS AND METHODS

### Drosophila Stocks and Transgenic Strains

All controls, stocks, and crosses were maintained on cornmeal medium (Archon Scientific, Durham, NC) at 25°C and 60% relative humidity with a 12/12 light/dark cycle in specialized incubators (Darwin Chambers, St. Louis, MO). Canton S. was used as the control line unless otherwise noted. All mutants and transgenes were maintained over larvally-selectable balancer chromosomes to enable identification. The following mutant alleles were used: *lrp4*^*dalek*^ (Mosca et al., 2017), *srpk79D*^*atc*^ (Johnson et al., 2009). The following UAS transgenes were used: *UAS-lrp4-HA* (Mosca et al., 2017), *UAS-mCD8-GFP* (T. Lee & Luo, 1999), *UAS-lrp4-RNAi* (108629, Vienna *Drosophila* Resource Center), *UAS-Dcr2* (Dietzl et al., 2007), *UAS-venus-SRPK79D-#28* (Johnson et al., 2009), *GMR90B08-GAL4* (referred to as *lrp4-GAL4*) was used to drive expression in cells expressing LRP4 (Pfeiffer et al., 2008). *C155-GAL4* (Lin & Goodman, 1994) was used to drive expression pan-neuronally. *OK6-GAL4* (Aberle et al., 2002) was used to drive expression in motor neurons. *Mhc-GAL4* (Schuster et al., 1996) or *DMef2-GAL4* (Lilly et al., 1995) was used to drive expression in all somatic muscles.

### Genotypes

See Table 1 for all genotypes.

### Construction of Fly Lines

An 3xHA-tag was knocked in to the endogenous *lrp4* locus to enable visualization of endogenous LRP4. We used CRISPR/ Cas9 genome editing (Gratz et al., 2015), with WellGenetics, Inc. (New Taipei City, Taiwan) to make a custom designed guide RNA and construct to introduce the 3xHA tag. We chose to tag LRP4 at the C-terminus, as a previous attempt generating a transgenic UAS line of LRP4 tagged at its C-terminus was successful (Mosca et al., 2017). Four lines were obtained and sequenced to confirm the presence of the 3xHA-tag, and each line was balanced over FM7a. We also used CRISPR to delete the *lrp4* coding sequence and generate an independent knock out allele *lrp4*^*Del*^ (Gratz et al., 2013). The deletion was confirmed by sequencing.

### Immunocytochemistry

Larvae were dissected and stained as previously described (Mosca & Schwarz, 2010; Restrepo et al., 2022). Larvae were raised in population cages (Genesee, no. 59-100) on grape juice plates supplemented with yeast paste. Wandering third instar larvae were dissected in Ca^2+^-free modified Drosophila saline (White et al., 2001). Larval fillets were fixed in 4% paraformaldehyde in 1x PBST for 20 min followed by three 20-minute washes in PBST, and 1 hour block in 5% normal goat serum. Samples were incubated in primary antibodies overnight, followed by three 10 min washes in PBST, and incubation in secondary antibodies for 2 hours at room temperature. The following primary antibodies were used: mouse anti-Brp (DSHB, cat. no. mAbnc82, 1:250) (Laissue et al., 1999), mouse anti-Dlg (DSHB, cat. no. mAb4F3, 1:500) (Parnas et al., 2001), rabbit anti-GluRIIC (custom, 1:2500) (Marrus et al., 2004), mouse anti-Futsch (DSHB, mAb22C10, 1:50) (Roos et al., 2000), rabbit anti-α Spectrin (custom, 1:1000) (Byers et al., 1989), rabbit anti-HA RM305 (RevMab Biosciences, Burlingame, CA, cat. no. 31-1190-00, 1:500). Alexa488-, Alexa647-, (Jackson ImmunoResearch, West Grove PA) and Alexa568-conjugated (ThermoFisher, Waltham, MA) secondary antibodies were used at 1:250. Cy3-or Alexa647-conjugated goat anti-HRP primary antibodies were used at 1:100 (Jackson ImmunoResearch, West Grove, PA). Samples processed for confocal imaging were mounted in Vectashield Antifade Mounting Medium (Vector Laboratories, Newark CA).

### Confocal Imaging and Image Processing

Confocal Z-stacks were acquired using a Zeiss LSM880 Laser Scanning Confocal microscope (Carl Zeiss, Oberlochen, Germany) with 40x 1.4 NA PlanApo or 63x 1.4 NA PlanApo oil objectives.

### STED Imaging and Deconvolution

For STED imaging, immunocytochemistry protocols were slightly adjusted to improve analysis of active zones (modified from Jetti et al., 2023). Following dissection in Ca^2+^ -free modified Drosophila saline, samples were fixed in 4% paraformaldehyde for 10 min. Following incubation in primary and secondary antibodies, samples were mounted on slides using SlowFade. All STED images were acquired as Z-stacks using a Leica TCS SP8 STED 3X microscope (Leica Microsystems, Wetzlar, Germany) with a 100x objective. For all STED imaging, mouse anti-Brp (DSHB, cat. no. mAbnc82, 1:250) primaries were used with Alexa488-conjugated secondaries. To deconvolve STED images, Z-stacks were first converted to stacked TIFF files using ImageJ and deconvolved using Nikon Elements software (Nikon Instruments, Melville, NY). 3D deconvolution was performed using the Landweber algorithm, with maximum 20 iterations. Deconvolved images were maximum intensity projected in ImageJ for analysis. Only type 1b boutons were analyzed.

### Electron Microscopy

Wandering third instar larvae were dissected as described above. Samples were fixed in 2.5% PFA, 5% glutaraldehyde, 0.06% picric acid in 0.1 M cacodylate buffer overnight, on ice. Following fixation, samples were incubated in 2% osmium tetroxide for one hour, on ice. Samples were then dehydrated in an ethanol series, rinsed in propylene oxide, and incubated in 50% propylene oxide / 50% resin overnight. Samples were then added to fresh resin for 4 hours and embedded in an incubator at 65°C for 2 days or until hard. The 6/7 muscle region was identified by taking 1 μm square sections, and bouton regions were located by taking 90 nm sections until muscle tissue was identified. All electron micrographs were acquired using a FEI Tecnai 12 120 keV digital TEM, fitted with a bottom-mounted AMT BioSprint 12 MPx CCD camera.

### Electrophysiology

Spontaneous and evoked postsynaptic potentials were recorded in voltage clamp mode from muscle 6 in male third-third instar larvae as previously described (Bruckner et al., 2017). Larvae were dissected in Ca^2+^ free hemolymph-like saline (HL3) which was replaced with saline containing 0.6 mM Ca^2+^ for recording. mEJPs were recorded for one minute, and 60 were averaged to obtain mEJP amplitude for each muscle before a stimulus was applied. EJPs were evoked in abdominal segment 3 and 4 by suctioning the cut end of the segmental nerve and applying stimulus at 0.5 Hz with stimulus amplitude adjusted to reliably evoke both Is and Ib nerve inputs. At least 25 consecutive EJPs were recorded for each cell and analyzed in pClamp to obtain mean amplitude. Quantal content was calculated for each cell as mean EJP amplitude divided by mean mEJP amplitude.

### Quantification of NMJ Synaptic Parameters

All NMJs were quantified from muscles 6/7 or muscle 4 on both the left and right sides, and comparisons were only made within larval segments. All phenotypes were also observed at other synapses regardless of muscle fiber or segment. Bouton number was counted in NMJs of muscles 6/7 by hand at segment A3, unless otherwise noted. Futsch loops and unbundled Futsch were counted by hand at terminals of muscles 6/7. Ghost boutons were quantified as HRP positive, Dlg-negative membrane protrusions with a visible connection to the NMJ terminal, as previously described (Restrepo et al., 2022). Synaptic spectrin fluorescence intensity was measured in ImageJ by drawing a region of interest surrounding the NMJ. Puncta density was determined in muscle 4 using the “Spots” function in Imaris software (Oxford Instruments, Abingdon, UK), with a spot size of 0.4 for Brp puncta, and 0.6 for GluRIIC puncta. Unapposed puncta were then counted by hand as either a Brp punctum lacking a corresponding GluRIIC, or a GluRIIC punctum lacking Brp.

Electron micrographs analyzed using ImageJ. Parameters for ultrastructural analysis were quantified as previously described (Mosca & Schwarz, 2010). SSR, PSD, and T-bar analysis was performed using ImageJ on boutons that were at least 1 μm in length and contained an active zone. Bouton area was calculated by tracing the perimeter of the bouton, and SSR area was calculated by tracing the perimeter of the bouton and the entire bouton + SSR and subtracting the area of the bouton. For SSR width, an arbitrary center point of the bouton was chosen, and 8 radii were drawn outward from the center at 45° angle intervals. The width of the SSR was measured at each line and averaged. For SSR complexity, 8 radii were drawn outward from the center at 45° angle intervals and the number of membranes crossing each line were counted by hand and averaged for each bouton.

Figures were constructed using ZEN 2.3 software (Carl Zeiss, Oberlochen, Germany), ImageJ (NIH, Bethesda, MD), Adobe Photoshop 2023, and Adobe Illustrator 2023 (Adobe Systems, San Jose, CA).

### Statistical Analysis

Statistical analysis was performed, and graphical representations prepared using Prism 9.5.1 (Graphpad Software, Inc., La Jolla, CA). Data is expressed as mean ± s.e.m. Normality was determined using a D’Agostino-Pearson test. Significance between two groups was determined using a two-tailed Student’s t-test. Significance among 3 or more groups was determined using one-way ANOVA with a Dunnett post-hoc test to a control group and a Bonferroni post-hoc test among all groups. Multiple comparisons were corrected for using a Tukey’s post-hoc test. For single comparisons between non-normally distributed data, a Mann Whitney U test was used. In each figure, unless otherwise noted, statistical significance is denoted in comparison to control genotypes.

---

**Figure S1.**
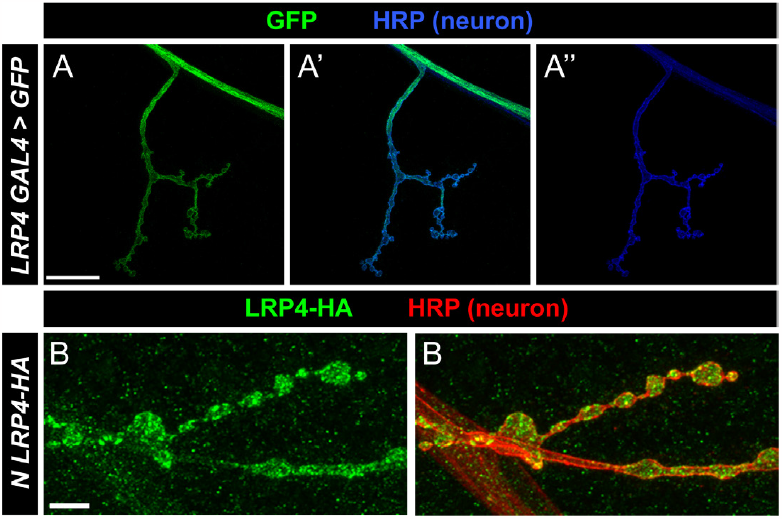
LRP4 is expressed in motor neurons and localizes in boutons at the NMJ. (**A**) Representative confocal image of an NMJ expressing *LRP4-GAL4* transgene and UAS-GFP and stained with antibodies to HRP (blue). GFP is observed in motor neurons. Scale bars = 20 μm. (B) Representative confocal image of an NMJ expressing HA-tagged LRP4 pan-neuronally using *C155-GAL4*, and stained with antibodies to HA (green) and HRP (red). LRP4-HA localizes to boutons. Scale bars = 5 μm.

**Figure S2.**
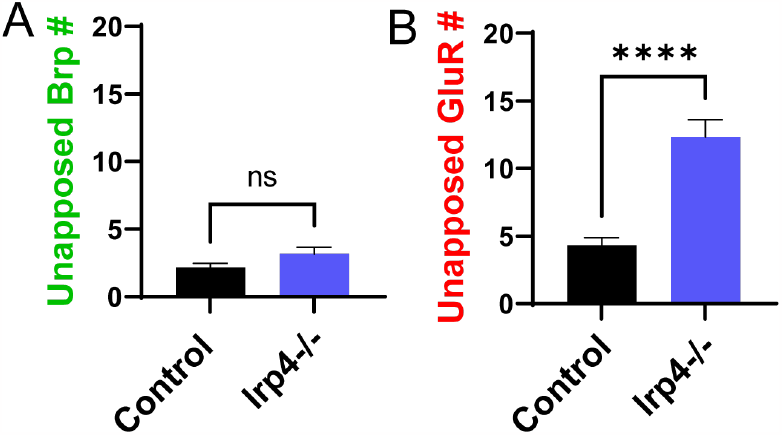
Loss of *lrp4* results in a significant increase in unapposed GluRIIC puncta. **(A)** Quantification of unapposed Brp puncta. Loss of *lrp4* does not significantly affect the number of Brp puncta lacking an apposite GluRIIC punctum. (**B**) Quantification of unapposed GluRIIC puncta. Loss of *lrp4* results in a significant increase in GluRIIC puncta lacking an apposite Brp punctum. For all experiments, data are shown as mean ± SEM. *p*\*\*\*\* < 0.001, *ns* = not significant. Significance was determined using a two-tailed Student’s t-test. n ≥ 12 NMJs, 6 larvae.

**Figure S3.**
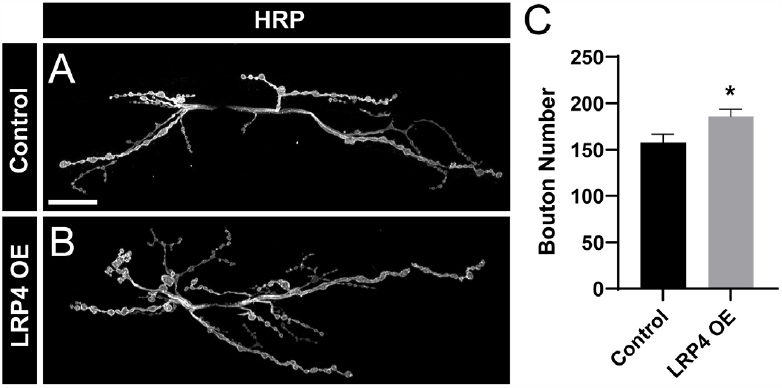
Overexpression of LRP4 increases bouton number. (**A-B**) Representative images of segment A2 NMJs from control (**A**) or following overexpression of LRP4 in neurons (**B**), stained with antibodies to HRP. Scale bars = 20 μm. (**C**) Quantification of bouton number. LRP4 overexpression results in a significant increase in bouton number. For all experiments, data are shown as mean ± SEM. *p*\* < 0.05. Significance was determined using a two-tailed Student’s t-test. n ≥ 12 NMJs, 7 larvae.

**Figure S4.**
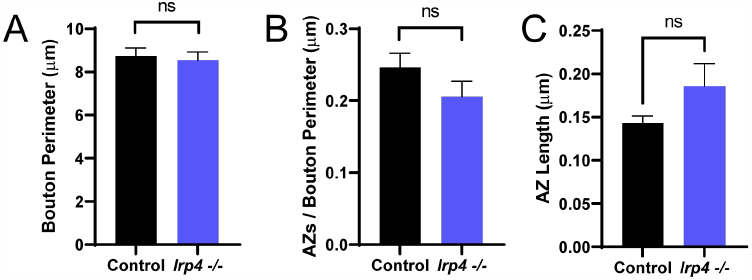
Loss of *lrp4* does not affect bouton perimeter or active zone number. (**A**) Quantification of bouton perimeter. Loss of *lrp4* does not affect bouton perimeter. (**B**) Quantification of AZs / bouton perimeter. No significant difference is observed between control and *lrp4* mutants. (**C**) Quantification of the length of the tabletop of the active zone T-bar. Loss of *lrp4* resulted in no significant change to AZ length. For all experiments, data are shown as mean ± SEM. ns = not significant. Significance was determined using a two-tailed Student’s t-test. n ≥ 44 boutons, 3 larvae.

**Figure S5.**
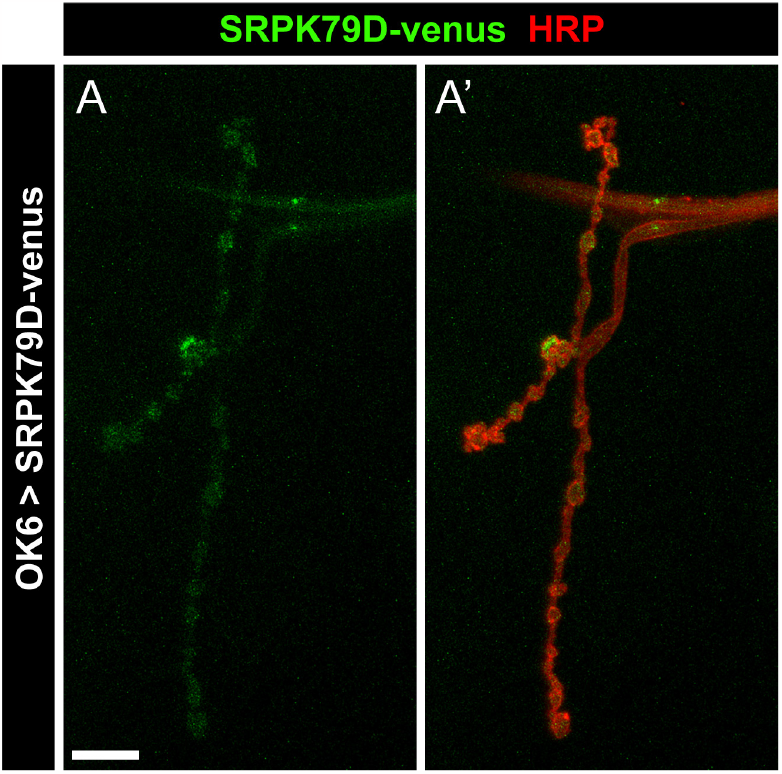
Tagged-SRPK79D localizes presynaptically. (**A**) Representative confocal image of an NMJ expressing venus-tagged SRPK79D (green) in motor neurons using OK6-GAL4 and stained with antibodies to HRP (red). SRPK79D localizes within the presynaptic compartment. Scale bars = 10 μm.

**Figure S6.**
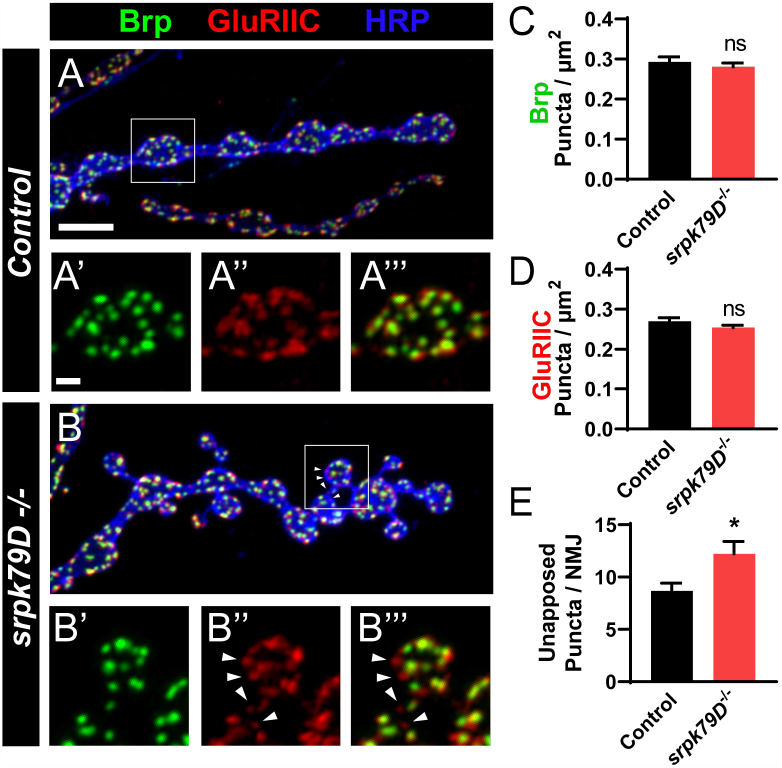
Active zone / receptor apposition is disrupted following loss of *srpk79D*. (**A-B**) Representative confocal images from control (**A**) and *srpk79D* mutant (**B**) NMJs stained with antibodies to Brp (green), GluRIIC (red), and HRP (blue). Arrowheads in (**B**) indicate unapposed puncta. Scale bars = 5 μm, 2 μm (insets). (**C**) Quantification of Brp density. (**D**) Quantification of GluRIIC density. (**E**) Quantification of the number of unapposed puncta per NMJ. Unapposed puncta number is significantly increased following loss of *srpk79D*. For all experiments, data are shown as mean ± SEM. *p*\* < 0.05, *ns* = not significant. Significance was determined using a two-tailed Student’s t-test. n ≥ 14 NMJs, 8 larvae.

**Figure S7.**
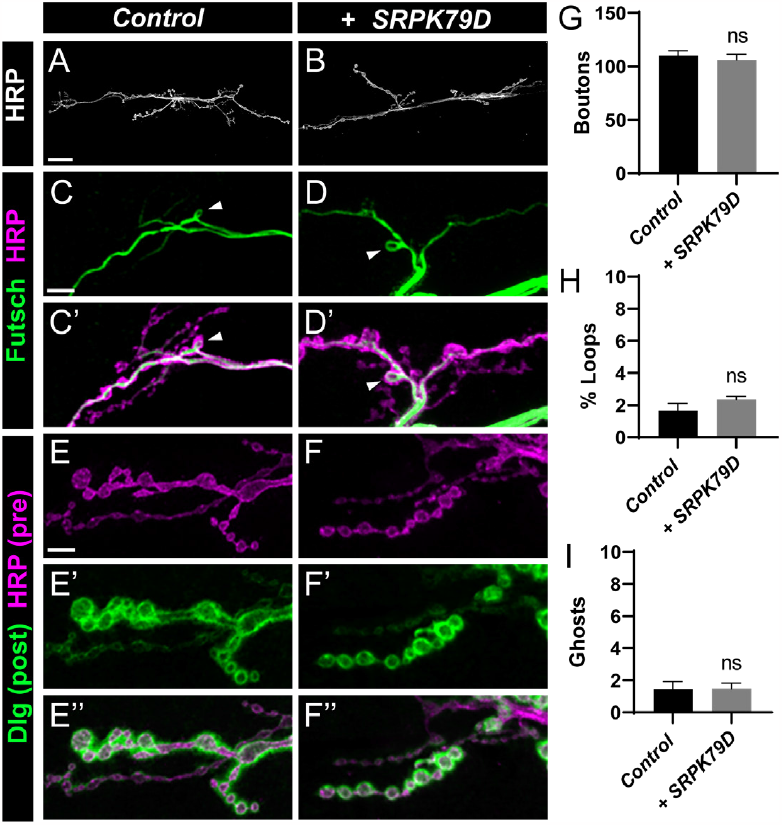
Overexpression of SRPK79D alone does not affect synapse growth or maturation. (**A-B**) Representative confocal images of NMJs from control (**A**) and with SRPK79D overexpression (**B**) stained with antibodies to HRP. Scale bars = 25 μm. (**C-D**) Representative confocal images of NMJs from control (**C**) and with SRPK79D overexpression (**D**) stained with antibodies to Futsch (green) and HRP (magenta). Scale bars = 5 μm. (**E-F**) Representative confocal images of NMJs from control (**C**) and with SRPK79D overexpression (**D**) stained with antibodies to Dlg (green) and HRP (magenta). Scale bars = 5 μm. (**G**) Quantification of bouton number, from experiments in (**A-B**). (**H**) Quantification of Futsch loops, from experiments in (**C-D**). (**I**) Quantification of ghost boutons, from experiments in (**E-F**). For all phenotypes, overexpression of SRPK79D resulted in no significant difference from control. For all experiments, data are shown as mean ± SEM. *ns* = not significant. Significance was determined using a two-tailed Student’s t-test. n ≥ 8 NMJs, 4 larvae.

